# Prophylactic Application of Tailocins Prevents Infection by *Pseudomonas syringae*

**DOI:** 10.1101/2020.08.31.276642

**Authors:** David A. Baltrus, Meara Clark, Kevin L. Hockett, Madison Mollico, Caitlin Smith, Savannah Weaver

## Abstract

Tailocins are phage-derived bacteriocins that demonstrate great potential as agricultural antimicrobials given their high killing efficiency and their precise strain-specific targeting ability. Our group has recently categorized and characterized tailocins produced by and tailocin sensitivities of the phytopathogen *Pseudomonas syringae*, and here we extend these experiments to test whether prophylactic tailocin application can prevent infection of *Nicotiana benthamiana* by *P. syringae pv. syringae* B728a. Specifically, we demonstrate that multiple strains can produce tailocins that prevent infection by strain B728a and engineer a deletion mutant to prove that tailocin targeting is responsible for this protective effect. Lastly, we provide evidence that heritable resistance mutations do not explain the minority of cases where tailocins fail to prevent infection. Our results extend previous reports of prophylactic use of tailocins against phytopathogens, and establish a model system with which to test and optimize tailocin application for prophylactic treatment to prevent phytopathogen infection.

## Introduction

Compared to the incredible advances that have occurred over the last century towards the clinical treatment of human and animal diseases, antibiotic treatments for prevention and control of infection of phytopathogens in plants have had relatively slower development and have been the subject of numerous debates over efficacy and effectiveness (Stockwell and Duffy 2012; McManus et al. 2002). Realistic options for the treatments of agricultural disease have further narrowed with heightened emphasis on the importance of one health initiatives as well as increasing recognition that broad spectrum antibiotics may have negative collateral effects on beneficial microbiota (Becattini et al. 2016; Robinson et al. 2016). With these ideas in mind we sought to develop and test for the ability of phage derived bacteriocins, which maintain a relatively precise and narrow spectrum killing activity against target strains, as a means to prevent infection by *Pseudomonas syringae*. We further describe this model system as a way to explore optimization of strain-specific prophylactic tailocin applications.

Recent efforts in the development of agricultural antimicrobials have turned towards developing specific and tailored treatments, such as the application of bacteriocins, as preventative measures for plant and animal disease (Buttimer et al. 2017; Rooney et al. 2020; Cotter et al. 2013; Behrens et al. 2017). Bacteriocins are a subset of antimicrobial compounds produced by various bacteria, which are largely thought to have a narrower spectrum of killing activity than more commonly used broad spectrum antibiotics (Chikindas et al. 2018; Riley and Wertz 2002). Increased specificity of bacteriocins occurs because these molecules must interact with dedicated receptor regions or proteins on target cells before antimicrobial activity is initiated (Behrens et al. 2017). Phage derived bacteriocins (also known as tailocins) are a subset of bacteriocins in which the phage tail structures have been coopted by bacteria through evolution to target and kill bacteria through binding and depolarization of their membranes (Patz et al. 2019). Tailocins have shown potential as a platform for the development of antimicrobials with high specificity against human and animal pathogens *in vitro* and *in vivo*, with additional demonstrated capability to engineer and expand target specificity through the incorporation of tail proteins from extant phage (Scholl et al. 2009; Williams et al. 2008; Ritchie et al. 2011). We have recently discovered and described a tailocin locus present in the phytopathogen *P. syringae*, in which the killing specificity of the tailocins is determined by interactions between receptor binding proteins and the Lipopolysaccharide (LPS) layer of target strains (Hockett et al. 2015; Kandel et al. 2020; Hockett et al. 2017). A recent report demonstrated that prophylactic tailocin application could prevent infection of tomatoes by *Xanthomonas (Príncipe et al. 2018)*, and we therefore tested whether these results could be extended to other phytopathogenic systems like *P. syringae*.

We have previously shown that tailocins from strains *Psy*Cit7 and USA011R can target strain *Psy*B728a, which can infect and cause disease in *Nicotiana benthamiana*. Here we establish that application of *P. syringae* tailocins (from strains *Psy*Cit7 and USA011R) prior to infection can prevent infection and disease in *Nicotiana benthamiana* caused by *P. syringae* strain PsyB728a. These results demonstrate that this protective ability is solely determined by the production and specificity of the tailocin molecules themselves and is conserved across different strains that produce tailocin molecules with similar killing spectra.

## Materials and Methods

### Bacterial Strains and Growth Conditions

*Psy*Cit7 was originally acquired from Steve Lindow and was described in XX. *P. syringae* strain USA011 was originally isolated by Cindy Morris (Morris et al. 2010). USA011R is a strain derived from USA011 by the Baltrus lab, isolated by plating out overnight cultures of USA011 on King’s Medium B (KB) rifampicin agar plates and selecting a single colony. This single colony was then picked to KB media with rifampicin and frozen at -80°C in 40% glycerol. This frozen isolate was used to create DBL1424. DBL1424 is a deletion mutant derived from USA011R, in which the R-type syringacin receptor binding protein (Rbp) and chaperone genes have been deleted using a method originally described in Baltrus et al. 2012 (Baltrus et al. 2012). For more details on creation of this deletion, please see 10.6084/m9.figshare.12814205. DBL1701 is a strain derived from DBL1424 in which deletion of the RBP and chaperone have been complemented and replaced *in cis*, please see 10.6084/m9.figshare.12814205 for additional details about complementation.

Typically, for all experiments, *P. syringae* isolates were grown at 27°C on King’s B (KB) agar and liquid media using rifampicin at 50 μg/ml. When necessary, cultures of both *P. syringae* and *Escherichia coli* were supplemented with antibiotics or sugars in the following concentrations: tetracycline at 10 μg/ml and 5% sucrose.

### Sequencing and Assembly of Bacterial Genomes

The same protocol was followed for culturing of each strain prior to DNA extraction for sequencing. A single colony arising from original frozen stocks streaked to KB agar media was picked to 2mL KB broth and grown overnight at 27°C in a shaking incubator at 220rpm. Genomic DNA used for Illumina sequencing and Nanopore sequencing was isolated from these 2mL overnight cultures via the Promega (Madison, WI) Wizard kit with the manufacturer’s protocols. RNAse A was added as per manufacturer’s protocols for all of the genomic isolations.

Genomic DNA from USA011R was sequenced by the Baltrus lab via an Oxford Nanopore MinION using a R9.4 flowcell, with 1ug of DNA prepared using the LSK-109 kit without shearing. Reads were called during sequencing using Guppy version 3.2.6 using a MinIT (ont-minit-release 19.10.3) for processing. Sequencing on the MinION generated ZZ reads for a total of ZZbp (∼ZZ coverage) of sequence with a read N50 of XXbp. Reads arising from Nanopore sequencing were used in conjunction with Illumina reads originally used to generate a draft sequence for this strain (Baltrus et al. 2014) using the hybrid assembler Unicycler and with default parameters. Log files for the assembly can be found at 10.6084/m9.figshare.12814205. This genome was annotated by NCBI’s PGAP pipeline (Tatusova et al. 2016).

For DBL1424 and DBL1701, DNA was sequenced by MiGS (Pittsburgh, PA) using an Illumina platform following their standard workflow for library preparation and read trimming. As described in (Baym et al. 2015), this workflow uses a Illumina tagmentation kit for library generation, followed by sequencing on a NextSeq 550 with 150 base pair (bp) paired-end reads. Trimmomatic (Bolger et al. 2014) was used for adaptor trimming using the default settings. This workflow generated a total of 1,555,032 paired reads and 413Mbp (∼68x coverage) of sequence for strain DBL1424 and 1,368,760 paired reads and 368Mbp (∼60x coverage) of sequence for strain DBL1701. These reads were fed into the Breseq pipeline (Deatherage and Barrick 2014) and analyzed against the complete genome sequence of USA011R with default parameters to confirm genotypes for strains DBL1424 and DBL1701.

### Tailocin Preparation and Quantification

Isolation of supernatants containing R-type syringacin molecules for strains USA011R, DBL1424, DBL1701, *Psy*B728a, and *P. aeruginosa* PAO1 and 1424 were completed as outlined in steps one and two of Hockett et al 2017 (Hockett and Baltrus 2017). Briefly, bacterial cultures were grown on KB agar plates for 48 hours at 27°C. A single colony was picked to 3 mL KB media and grown overnight shaking at 27°C. The strains were then back diluted 1:100 and grown for 3-4 hours, then 2uL mitomycin C was added with a final concentration of 0.5 μg/ml in 3mL. The culture was then grown overnight and sterilized with chloroform the next day, and stored at 4°C until further use. The quantification of the tailocins was completed following the methods outlined in (Haag and Vidaver 1974).

### Plant Infections

Prior to infection experiments involving tailocins from *Psy*Cit7, individual seeds of *Nicotiana benthamiana* were placed into individual peat pellets (Jiffy, Lorain OH) and germinated and grown in the laboratory window using natural light. Plants were maintained in the window for 3-4 weeks in domed flats at which point they were used for infection experiments. Prior to infection involving tailocins from USA011R strains, *N. benthamiana* plants in domed flats were germinated and grown for 2-3 weeks in a growth chamber under 18L/6d scheme and at 65% humidity at which point they were moved to a laboratory window for infection experiments.

For infections, a small amount of *Psy*B728a was picked from a KB agar plate and grown overnight in KB media, pelleted and washed twice with 10Mm MgCl_2_, and then resuspended in an inoculation solution of 10Mm MgCl_2_ and silwet (40uL per 200mL) with bacteria at an OD600 of 0.05. For tailocin treatments, bacterial supernatants for each treatment were painted onto individual plant leaves using a sterile Qtip until the entire plant was covered. Plants treated with supernatants were left undomed for 1 hour, at which point they were dipped into the inoculum containing bacterial strain *Psy*B728a. Plants were then maintained in the laboratory window in domed flats until bacteria were sampled. At least 2 plants were infected for each treatment for each replicate (although the far majority of treatments consisted of 4 or greater plants per replicate), and each set of experiments consisted of a total of three replicates spread out over multiple weeks.

At 3 days post inoculation, the most diseased leaf from each plant was harvested into 500mL 10Mm MgCl2. These infected leaves were macerated with small beads using a MP FastPrep-24 for 2 cycles at 20s per cycle. A dilution series from each of these samples was then plated out on KB media containing rifampicin and bacteria were enumerated after 3 days.

## Results

### Application of tailocins from *P. syringae* strain Cit7 Protect *N. benthamiana* from infection by *Psy*B728a

Strain *Psy*Cit7 produces a tailocin that is active against *P. syringae* strain *Psy*B728a, while the tailocin produced by *Psy*B728a has no measurable activity against itself (Baltrus et al. 2019). Our *in planta* experiments also include additional negative controls which consist of preparations of R-type pyocins from *Pseudomonas aeruginosa*, which do not have measurable activity against strain *Psy*B728a (data not shown), as well as a treatment with no supernatants applied to the plants. As one can see in Fig. 1, across 3 replicate trials, supernatant preparations from strain *Psy*Cit7 containing a tailocin against *Psy*B728a provide extensive protection against infection of *N. benthamina* if applied prophylactically before plants were inoculated with the pathogen compared to the no tailocin controls and the other treatments. This difference is statistically supported by a Kruskal-Wallis test (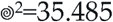,df=3, p<0.0001) followed by pairwise Wilcox tests between each treatment (*Psy*Cit7 vs. No Tailocin, p<0.0001). These results stand in direct contrast to those found in the two other treatments (supernatants from *Psy*B728a vs. No tailocin, p=0.18) and *P. aeruginosa* PAO1 vs. No Tailocin p=0.53), which did not provide any additional protection from infection compared to the no tailocin control. Overall, the most striking result is that no cells of *Psy*B728a could be recovered by plating from plants treated with supernatants containing Cit7 tailocins in 11/16 replicates.

**Figure 1:**
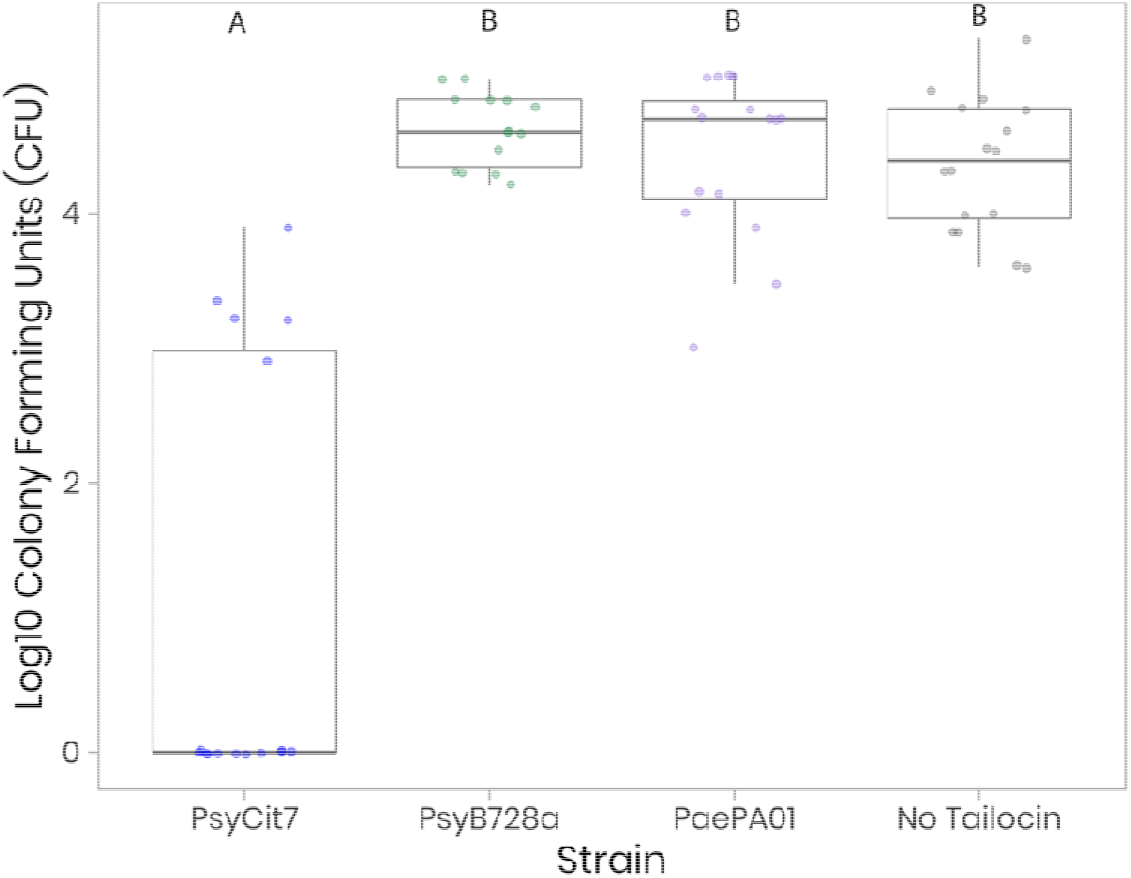
Supernatants Containing Tailocins from Strain *Psy*Cit7 Protect *N. benthamiana* from Infection by *P. syringae Psy*B728a. Shown on the Y-axis is the amount of strain *Psy*B728a recovered from an infected *N. benthamiana* leaf at 3 days post infection. Plants were pretreated with supernatants from a variety of strains (or no supernatant at all). Supernatants were produced by a variety of strains shown on the X-axis: *P. syringae Psy*Cit7 (which produces a tailocin that can target strain *Psy*B728a); *P. syringae Psy*B728a (which produces a tailocin but does not target itself); *P. aeruginosa* PA01 (which produces an R-type pyocin which does not target *Psy*B728a); and no supernatant applied. Data was gathered across three different experiments with at least 2 (and most often 4) replicates per experiment per strain. Groups tthat are significantly different (p<0.01) are differentiated by letters according to results of pairwise Wilcox tests with correction for multiple testing.

### Deletion of the Tailocin Receptor Binding Protein and Chaperone from USA011R Eliminates Killing Activity of Strain USA011R Against *Psy*B728a

In order to genetically test whether the production of active tailocins was required for protection of *N. benthamiana* from *Psy*B728a, we created a deletion mutant in which the tailocin receptor binding protein and chaperone were deleted from strain USA011R. We have previously shown that strain USA011R produces a tailocin that can specifically target *Psy*B728a (Baltrus et al. 2019). Whole genome sequencing of this strain confirmed that deletion was created as intended in strain DBL1424 (see Breseq results in 10.6084/m9.figshare.12814205). We further sought to complement the deletion within strain DBL1424 by replacing the deleted region *in cis* with DNA that nearly matched the original sequence from USA011R. Whole genome sequencing of DBL1701 confirmed that this region was successfully replaced with the exception that there is one silent single nucleotide polymorphism in the receptor binding protein compared to USA011R (see Breseq results in 10.6084/m9.figshare.12814205). As shown in Fig. 2, deletion of the receptor binding protein and chaperone in strain DBL1424 eliminates killing activity by this strain against *Psy*B728a. Furthermore, killing activity is restored when genes that encode production of these proteins are replaced *in cis* in strain DBL1701.

**Figure 2:**
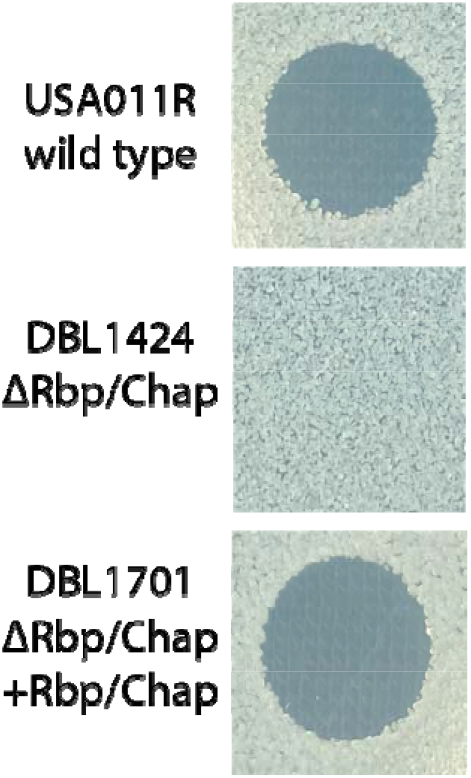
Deletion of the Receptor Binding Protein and Chaperone from Strain USA011R Eliminates Tailocin Killing of Strain *Psy*B728a. An overlay experiment, as per XX, was carried out in which tailocins were produced by strains USA011R, DBL1424, and DBL1701. A clearing zone indicates that strain *Psy*B728a is killed by tailocins from both USA011R and DBL1701, and a lack thereof demonstrates that this strain is not killed by tailocins from strain DBL1424.

### Protection of *N. benthamiana* by *P. syringae* strain USA011R is Dependent on Functional Tailocin Production

In order to clearly demonstrate that active tailocins were the factor limiting infection of plants by *Psy*B7278a in our previous experiments, we repeated the tailocin protection assays using strain USA011R which possesses an R-type syringacin that is capable of targeting and killing *Psy*B728a. In this second set of trials, we included supernatants prepared from a mutant of strain DBL1424 in which genes encoding the tailocin receptor binding protein and its chaperone were cleanly deleted. Lastly, we included supernatants from strain DBL1701, in which the genes cleanly deleted from strain DBL1424 were replaced *in cis* such that tailocin production was phenotypically complemented. As one can see in Figure 3A, preparations containing tailocins from strains USA011R and DBL1701 were able to protect *N. benthamiana* from infection by *Psy*B728a when applied as a prophylactic. This difference is statistically supported by a Kruskal-Wallis test (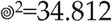,df=4, p<0.0001) followed by pairwise Wilcox tests between each treatment (USA011R vs DBL1424, p=0.00021; DBL1701 vs. DBL1424, p=0.00057). However, supernatant preparations from strain DBL1424 (which are essentially identical to those from USA011R and DBL1701 except that the tailocin structures lack receptor binding proteins) showed no significant difference in the ability to protect plants from *Psy*B278a infection from either the *Psy*B728a supernatant preparation or the no tailocin treatment. As with *Psy*Cit7, no viable cells were recovered from a majority of plants treated with either USA011R supernatants (9/12) or DBL1701 supernatants (8/10). These results were echoed in an experiment where plants were allowed to grow to 11dpi (Fig. 3C).

**Figure 3:**
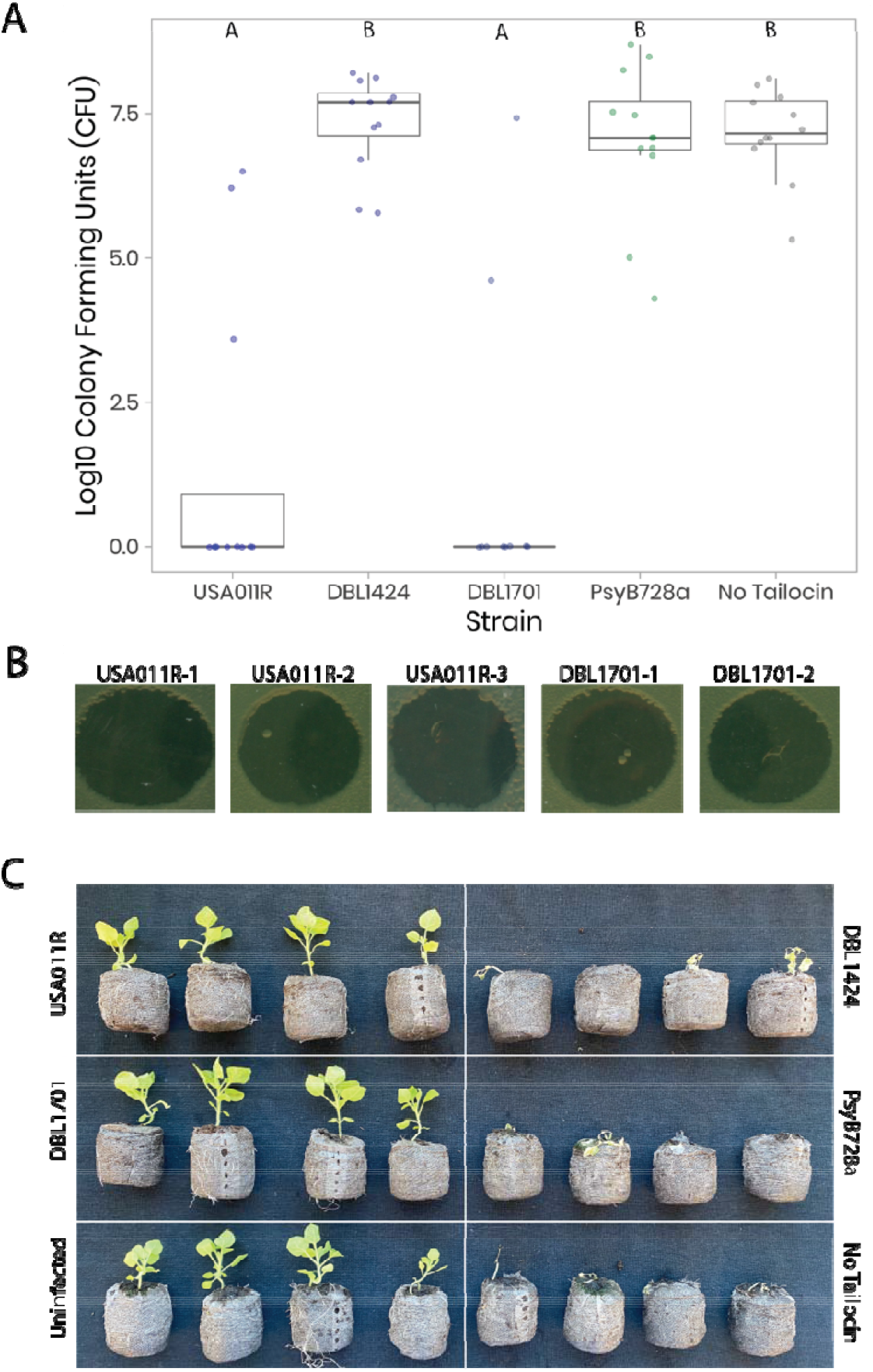
Tailocins from Strain USA011R Protect *N. benthamiana* from Infection by *P. syringae Psy*B728a. **A)** Shown on the Y-axis is the amount of strain *Psy*B728a recovered from an infected *N. benthamiana* leaf at 3 days post infection. Plants were pretreated with supernatants from a variety of strains (or no supernatant at all) derived from *P. syringae* USA011R. DBL1424 is a deletion mutant in which the Receptor Binding Protein (Rbp) of the tailocin has been deleted. DBL1701 is a derivative of strain DBL1424 in which the Rbp and chaperone from strain USA011R was replaced for complementation of strain DBL1424 *in trans*. Data was gathered across three different experiments with at least 2 (and most often 4) replicates per experiment per strain. Groups that are significantly different (p<0.01) are differentiated by letters according to results of pairwise Wilcox tests with correction for multiple testing. **B)** Single colonies were picked from plants infected despite application of preventative tailocins, and tested for sensitivity to these tailocins. All three colonies arising from plant infection after application of USA011R tailocins and both colonies arising from plant infection after DBL1701 tailocins maintain sensitivity to killing by USA011R tailocins. C) An additional trial was carried out in which plants were pretreated with supernatants containing tailocins for the same strains as in A) except that this experiment also included a no infection control. In this experiment, plants were photographed at 11dpi and there were 4 replicate plants per treatment.

As with the Cit7 protection assays above, there was growth of strain *Psy*B728a during a limited number of replicates using supernatant preparations from either USA011R (3/12) or DBL1701 (2/10 infections). In this case, a single colony of strain *Psy*B728a was picked from most diluted sample arising from each infected plant and tested for resistance to tailocins from USA011R. Every one of these tested colonies maintained tailocin sensitivity (Fig 3B).

## Discussion

Given their high target specificity and efficient killing capability, phage derived bacteriocins could provide important prophylactic treatments for agricultural crops against a suite of phytopathogens (Mills et al. 2017). Indeed, previous reports strongly suggested that application of tailocins from Pseudomonas could prevent infection of tomatoes by a *Xanthomonas* strain (Príncipe et al. 2018). As an added benefit, resistance mutations arising against tailocins have also been shown to severely affect the virulence of plant pathogens and may even sensitize strains to killing by alternative antimicrobials, so that the evolution of tailocin resistance under natural conditions may be inhibited by these tradeoffs (Kandel et al. 2020; Hockett et al. 2017). Lastly, since tailocins are non-replicating structures composed only of protein, it is possible that they could be more durable than phage treatments under the harsh conditions of agricultural fields and would not suffer from worries associated with uncontrolled release of phage (Meaden and Koskella 2013). Therefore, our goal with this manuscript was to extend these previously reported results to a new pathosystem and to establish a genetically controlled model system with which to begin to optimize tailocin application in which to see protective effects and which could be used to explore conditions to ensure efficient plant protection under a variety of conditions.

We demonstrate that tailocin production by two different strains (*Psy*Cit7 and USA011R) can effectively block infection of *N. benthamiana* by the phytopathogen *P. syringae* pv. *syringae* B728a (Figs. 1 and 3). These treatments proved highly effective, and in a far majority of cases there were no cells of *Psy*B728a recovered from plants even though these plants were dipped in inoculum containing 10^7^ CFU/mL. In contrast, there was vigorous growth and infection of plants by *Psy*B728a under control treatments where no supernatants were added to the plants prior to infection. These results are especially clear at 11dpi (Figure 3B), where the tailocin treated plants are growing quite vigorously while the plants without protection of tailocins are dead or nearly so. Our experiments also demonstrate that supernatants prepared from alternative strains, which produce tailocins that do not target *Psy*B728a, do not provide any enhanced protection of plants than the no tailocin control.

Although our initial experiments were highly suggestive (Fig. 1), the possibility remained that supernatants from strain *Psy*Cit7 contained additional (non-tailocin) molecules compared to those from either *Psy*B728a or *P. aeruginosa* that could potentially protect plants from infection or which could stimulate plant defenses prior to infection. To address this critique, we generated a mutant strain of USA011R (referred to here as DBL1424) in which the tailocin receptor binding protein and its chaperone were deleted from the genome as well as a strain (DBL1701) in which this deleted region was replaced with sequence nearly identical to the region from strain USA011R. Deletion of these two genes in strain DBL1424 eliminates tailocin killing activity against strain *Psy*B728a, and this activity is phenotypically complemented in strain DBL1701 (Fig. 2). Supernatants produced by strain DBL1424 are nearly identical to those produced by strain USA011R, except that the tailocins in supernatants from strain DBL1424 cannot bind to target cells and thus have no killing activity. Experiments *in planta* using both of these strains (Fig. 3) clearly demonstrate that active tailocins are necessary to provide protection to plants against infection by strain *Psy*B728a.

In a small number of plants, tailocin treatments were ineffective in preventing infection by *Psy*B728a. We currently do not have an explanation for these results, but tested whether these infections were enabled through the evolution of genetic resistance against tailocins by isolating colonies arising from these infected plants and testing for tailocin resistance using overlay assays. In no case did we see genetic resistance against tailocins from USA011R when these colonies were restested. It remains a possibility that these rare instances of tailocin evasion were the product of persister like phenotypes against tailocins which were recently reported (Kandel et al. 2020). It could also be that a subset of cells periodically switches between tailocin resistance and sensitivity (perhaps through LPS modification (Simpson and Trent 2019)), and that this switch resets quickly enough to tailocin sensitivity when strains are grown under conditions for overlay experiments. Lastly, it may simply be that our crude application of tailocins to leaves was suboptimal in some cases and that with future experiments we could optimize tailocin application to ensure complete protection.

Although we’ve demonstrated the ability of tailocins to serve as a source of prophylatic protection against infection, many bacterial diseases are not treated prophylactically for economic reasons, especially in vegetable crops. Future studies will assess whether application of tailocins is able to prevent spread of the pathogen within a field after a focal disease outbreak has occurred. It will also be of use to assess whether post-disease application of tailocins will be able to prevent formation of secondary infections on the same leaf. Lastly, we note that these and other studies point towards potentially engineering tailocins to be produced by plants as an additional layer of resistance against bacterial phytopathogens.

In sum, we report that application of phage derived bacteriocins to the leaves of *N. benthamiana* under the conditions described herein can reliably provide complete protection against infection by *P. syringae* strain B728a. These results support previous reports describing how application of tailocins to plants could prevent infection by phytopathogens. Our experiments expand on these previous reports by including a variety of different phenotypic and genetic controls which enable the clear attribution of causality to tailocins for these protective effects. We look forward to building on this system to optimize tailocin treatments to provide complete plant protection while also exploring the limits of tailocin protection of many different plant hosts against a wide range of phytopathogens.

## Data Availability

Datasets underlying the results shown in Figures 1 and 3A, as well as the R commands used to create the figures, can be accessed at <u>doi:10.5281/zenodo.3987443</u>. A roadmap for creating deletion in strain DBL1424 as well as for creating the complementation strain DBL1701 can be found at 10.6084/m9.figshare.12814205. Unmodified pictures used to create Figures 2 and 3B can be found at 10.6084/m9.figshare.12814205. A complete genome sequence for strain USA011 is found at NCBI at accession GCA_000452525.4, and raw sequencing read files used to assemble the genome of USA011R as well as to confirm genotypes of DBL1424 and DBL1701 can be found in the SRA at SRR12516783 and SRR12516782 respectively..

## Acknowledgements

This work was partially supported by grants from the US Department of Agriculture (USDA) NIFA 2016-67014-24805 and National Science Foundation (NSF) IOS 1856556 to DAB. Partial support also came from the UBRP program and the BMCB training grant at the University of Arizona.

## Notes

https://doi.org/10.6084/m9.figshare.12814205

https://doi.org/10.5281/zenodo.3987443

### Competing Interest Statement

The authors have declared no competing interest.

